# Far-UVC light: A new tool to control the spread of airborne-mediated microbial diseases

**DOI:** 10.1101/240408

**Authors:** David Welch, Manuela Buonanno, Veljko Grilj, Igor Shuryak, Connor Crickmore, Alan W. Bigelow, Gerhard Randers-Pehrson, Gary W. Johnson, David J. Brenner

## Abstract

Airborne-mediated microbial diseases such as influenza and tuberculosis represent major public health challenges. A direct approach to prevent airborne transmission is inactivation of airborne pathogens, and the airborne antimicrobial potential of UVC ultraviolet light has long been established; however, its widespread use in public settings is limited because conventional UVC light sources are both carcinogenic and cataractogenic. By contrast, we have previously shown that far-UVC light (207–222 nm) efficiently kills bacteria without harm to exposed mammalian skin. This is because, due to its strong absorbance in biological materials, far-UVC light cannot penetrate even the outer (non living) layers of human skin or eye; however, because bacteria and viruses are of micrometer or smaller dimensions, far-UVC can penetrate and inactivate them. We show for the first time that far-UVC efficiently kills airborne aerosolized viruses, a very low dose of 2 mJ/cm^2^ of 222-nm light inactivating >95% of aerosolized H1N1 influenza virus. Continuous very low dose-rate far-UVC light in indoor public locations is a promising, safe and inexpensive tool to reduce the spread of airborne-mediated microbial diseases.

---

Airborne-mediated microbial diseases represent one of the major challenges to worldwide public health^1^. Common examples are influenza^2^, appearing in seasonal^3^ and pandemic^4^ forms, and bacterially-based airborne-mediated diseases such as tuberculosis^5^, increasingly emerging in multi-drug resistant form.

A direct approach to prevent the transmission of airborne-mediated disease is inactivation of the corresponding airborne pathogens, and in fact the airborne antimicrobial efficacy of ultraviolet (UV) light has long been established^6–8^. Germicidal UV light can also efficiently kill both drug-sensitive and multi-drug-resistant bacteria^9^, as well differing strains of viruses^10^. However, the widespread use of germicidal ultraviolet light in public settings has been very limited because conventional UVC light sources are a human health hazard, being both carcinogenic and cataractogenic^11,12^.

By contrast, we have earlier shown that far-UVC light generated by filtered excimer lamps emitting in the 207 to 222 nm wavelength range, efficiently kills drug-resistant bacteria, without apparent harm to exposed mammalian skin^13–15^. The biophysical reason is that, due to its strong absorbance in biological materials, far-UVC light does not have sufficient range to penetrate through even the outer layer (stratum corneum) on the surface of human skin, nor the outer tear layer on the outer surface of the eye, neither of which contain living cells; however, because bacteria and viruses are typically of micron or smaller dimensions, far-UVC light can still efficiently traverse and inactivate _them_^13–15^.

The earlier studies on the germicidal efficacy of far UVC light^13,15–18^ were performed exposing bacteria irradiated on a surface or in suspension. In that a major pathway for the spread of influenza A is aerosol transmission^3^, we investigate for the first time the efficacy of far-UVC 222-nm light for inactivating airborne viruses carried by aerosols – with the goal of providing a potentially safe alternative to conventional 254-nm germicidal lamps to inactivate airborne microbes.

## Results

### Virus inactivation

Fig. 1 shows representative fluorescent 40x images of mammalian epithelial cells incubated with airborne viruses that had been exposed in aerosolized form to far-UVC doses (0, 0.8, 1.3 or 2.0 mJ/cm^2^) generated by filtered 222-nm excimer lamps. Blue fluorescence was used to identify the total number of cells in a particular field of view, while green fluorescence indicated the integration of live influenza A (H1N1) viruses into the cells. Results from the zero-dose control studies (Fig. 1, top left) confirmed that the aerosol irradiation chamber efficiently transmitting the aerosolized viruses through the system, after which the live virus efficiently infected the test mammalian epithelial cells.

**Figure 1.**
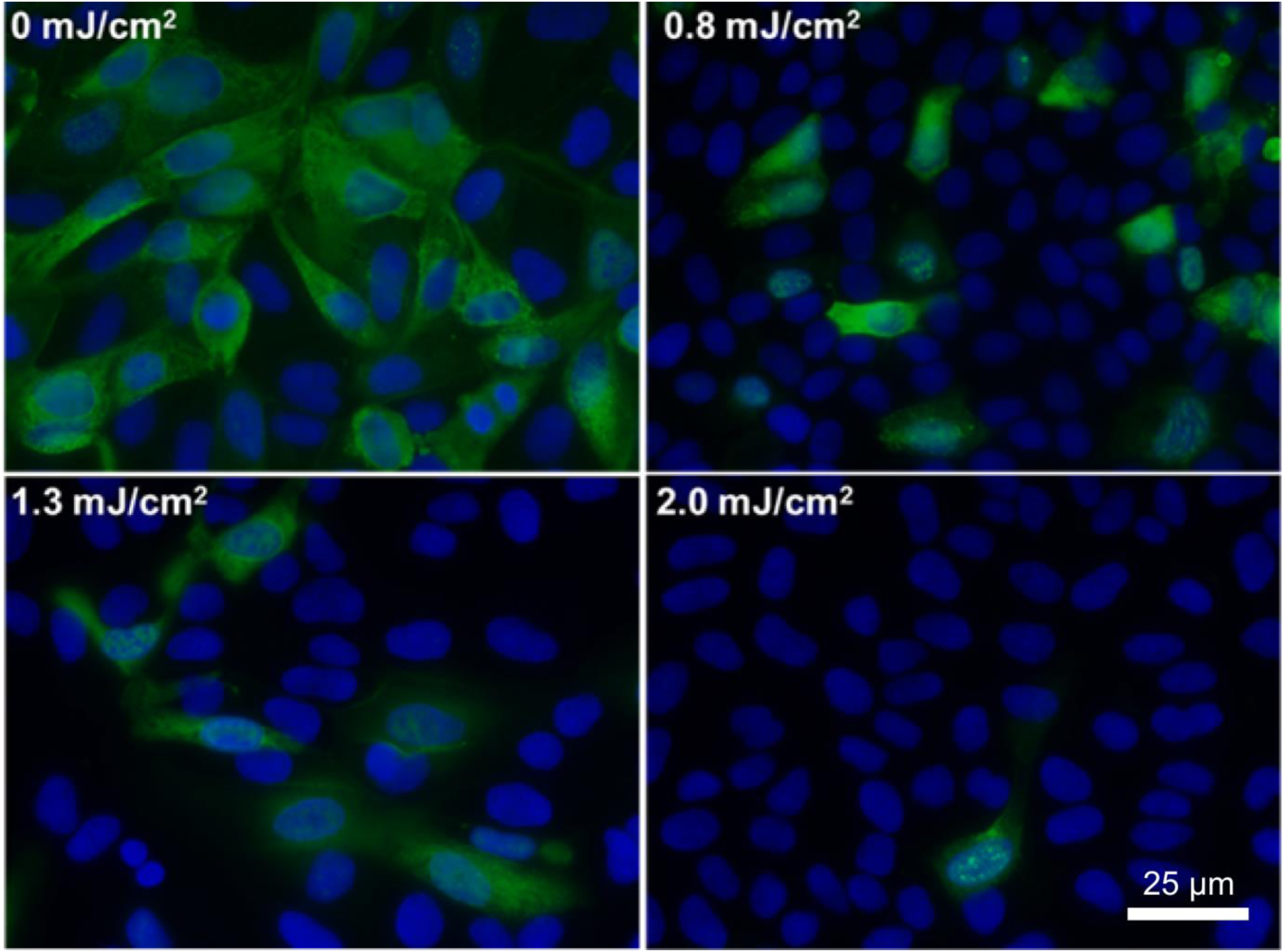
Antiviral efficacy of different low doses of 222-nm far-UVC light. Typical fluorescent images of MDCK epithelial cells infected with influenza A virus (H1N1). The viruses were exposed in aerosolized form in the irradiation chamber to doses of 0, 0.8, 1.3 or 2.0 mJ/cm^2^ of 222-nm far-UVC light. Infected cells fluoresce green (blue = nuclear stain DAPI; green= Alexa Fluor-488 conjugated to anti-influenza A antibody). Images were acquired with a 40x objective.

Fig. 2 shows the surviving fraction, as a function of the incident 222-nm far-UVC dose, of exposed H1N1 aerosolized viruses, as measured by the number of focus forming units in incubated epithelial cells relative to unexposed controls. Linear regressions (see below) showed that the survival results followed a classical exponential UV disinfection model with rate constant *k*=1.8 cm^2^/mJ (95% confidence intervals 1.5–2.1 cm^2^/mJ). The overall model fit was good, with a coefficient of determination, R^2^= 0.95, which suggests that most of the variability in virus survival was explained by the exponential model. The rate constant of 1.8 cm^2^/mJ corresponds to an inactivation cross-section (dose required to kill 95% of the exposed viruses) of D_95_ = 1.6 mJ/cm^2^ (95% confidence intervals 1.4–1.9 mJ/cm^2^).

**Figure 2.**
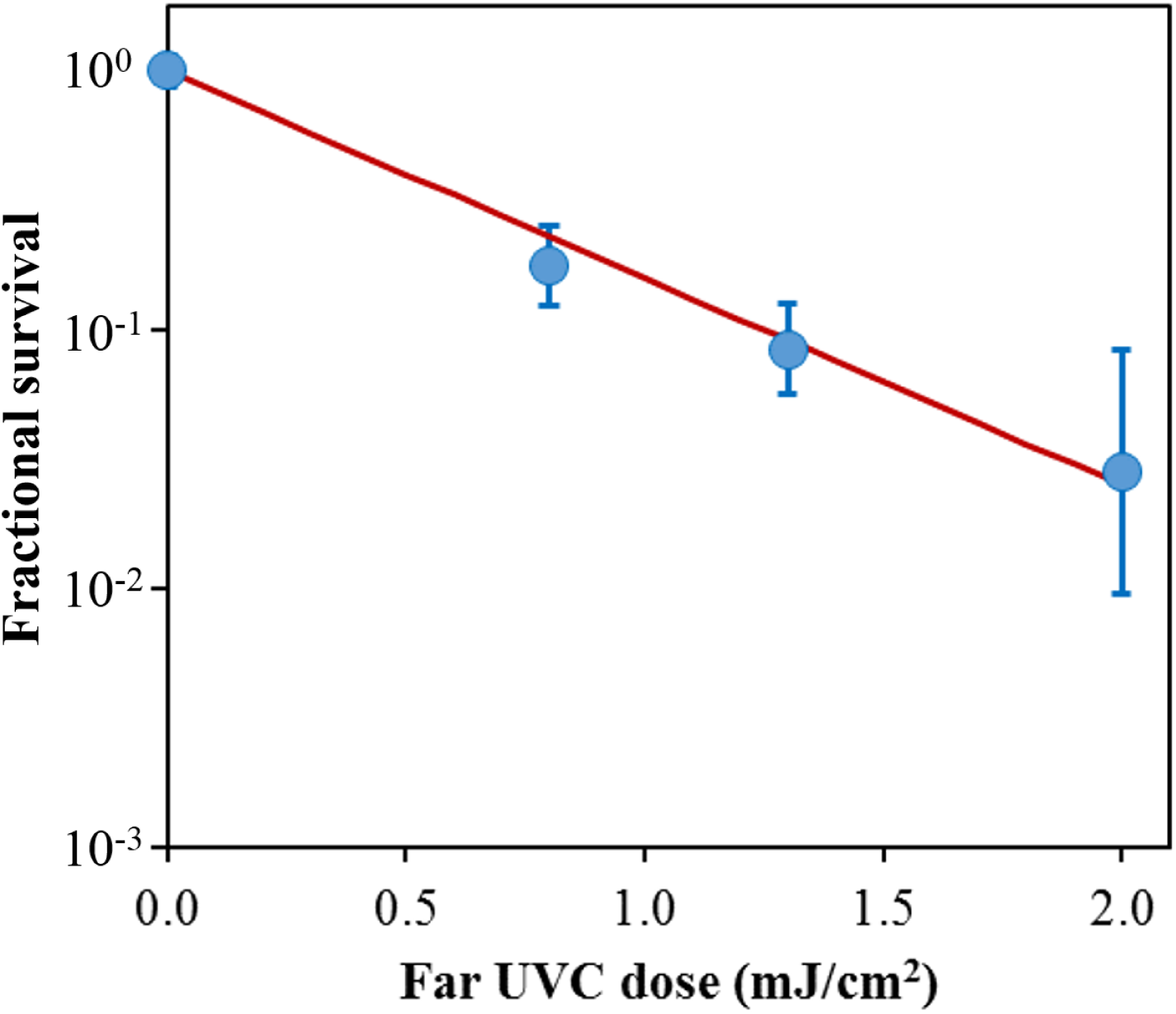
Quantification of the antiviral efficacy of 222-nm far-UVC light. Fractional survival, FFU_uv_/FFU_controls_, is plotted as a function of the 222-nm far-UVC dose. Means and standard deviations refer to triplicate repeat studies and the line represents the best-fit regression to Eqn 1 (see text).

## Discussion

We have developed an approach to UV-based sterilization using single-wavelength far-UVC light generated by filtered excilamps, which selectively inactivate microorganisms, but does not produce biological damage to exposed mammalian cells and tissues^13–15^. The approach is based on biophysical principles in that far-UVC light can traverse and therefore kill bacteria and viruses which are typically micrometer dimensions or smaller, whereas due to its strong absorbance in biological materials, far-UVC light cannot penetrate even the outer dead-cell layers of human skin, nor the outer tear layer on the surface of the eye.

Here we applied this approach to test the efficacy of the 222-nm far-UVC light to kill influenza A virus (H1N1) carried by aerosols in a benchtop aerosol UV irradiation chamber, which generated aerosol droplets of sizes similar to those generated by human coughing and breathing. Aerosolized viruses flowing through the irradiation chamber were exposed to UVC emitting lamps placed in front of the chamber window.

As shown in Fig. 2, killing of influenza A virus (H1N1) by 222-nm far-UVC light follows a typical exponential disinfection model, with an inactivation cross-section of D_95_ = 1.6 mJ/cm^2^ (95% CI: 1.4–1.9). For comparison, using a similar experimental arrangement, but using a conventional 254 nm germicidal UVC lamp, McDevitt *et al.^19^* found a D95 value of 1.1 mJ/cm^2^ (95% CI: 1.0–1.2) for H1N1 virus. Thus as we^13,15^ and others ^16–18^ reported in earlier studies for bacterial inactivation, 222-nm far-UVC light and 254-nm broad-spectrum germicidal light are quite similar in their efficiencies for viral inactivation, the comparatively small differences presumably reflecting differences in nucleic acid absorbance. However as discussed above, based on biophysical considerations and in contrast to the known human health safety issues associated with conventional germicidal 254-nm broad-spectrum UVC light, far-UVC light does not appear to be cytotoxic to exposed human cells and tissues *in vitro* or *in* vivo^13–15^.

If these results are confirmed in other scenarios, it follows that the use of overhead low-level far-UVC light in public locations may represent a safe and efficient methodology for limiting the transmission and spread of airborne-mediated microbial diseases such as influenza and tuberculosis. In fact the potential use of ultraviolet light for airborne disinfection is by no means new, and was first demonstrated more than 80 years ago^8,20^. As applied more recently, airborne ultraviolet germicidal irradiation (UVGI) utilizes conventional germicidal UVC light in the upper part of the room, with louvers to prevent direct exposure of potentially occupied room areas^21^. This results in blocking more than 95% of the UV radiation exiting the UVGI fixture, with substantial decrease in effectiveness^22^. By contrast, use of low-level far-UVC fixtures, which are potentially safe for human exposure, could provide the desired antimicrobial benefits without the accompanying human health concerns of conventional germicidal lamp UVGI.

A key advantage of the UVC based approach, which is in clear contrast to vaccination approaches, is that UVC light is likely to be effective against all airborne microbes. For example, while there will almost certainly be variations in UVC inactivation efficiency as different influenza strains appear, they are unlikely to be large^7,10^. Likewise, as multi-drug-resistant variants of bacteria emerge, their UVC inactivation efficiencies are also unlikely to change greatly^9^.

Finally it is of course by no means the case that all microbes are harmful, and it is well established that the human microbiome is essential to human health^23^. With the exception of a subset of the skin microbiome, all the human microbiome would be entirely shielded from far-UVC light due to its very short range; in fact even within the skin biome only those biota on the skin surface^24^ would be potentially affected, which are of course the same biota that are potentially removed by hand sanitizers^25^.

In conclusion, we have shown for the first time that very low doses of far-UVC light efficiently kill airborne viruses carried by aerosols. For example, a very low dose of 2 mJ/cm^2^ of 222-nm light inactivates >95% of airborne H1N1 virus. Our results indicate that far-UVC light is a powerful and inexpensive approach for prevention and reduction of airborne viral infections without the human health hazards inherent with conventional germicidal UVC lamps. If these results are confirmed in other scenarios, it follows that the use of overhead very low level far-UVC light in public locations may represent a safe and efficient methodology for limiting the transmission and spread of airborne-mediated microbial diseases. Public locations such as hospitals, doctors’ offices, schools, airports and airplanes might be considered here. This approach may help limit seasonal influenza epidemics, transmission of tuberculosis, as well as major pandemics.

## Methods

### Far-UVC lamps

We used a bank of three excimer lamps containing a Kr-Cl gas mixture that predominantly emits at 222 nm ^26,27^. The exit window of each lamp was covered with a custom bandpass filter designed to remove all but the dominant emission wavelength as previously described^15^. Each bandpass filter (Omega Optical, Brattleboro, VT) had a center wavelength of 222 nm and a full width at half maximum (FWHM) of 25 nm and enables >20% transmission at 222 nm. A UV spectrometer (SPM-002-BT64, Photon Control, BC, Canada) with a sensitivity range between 190 nm and 400 nm was utilized to verify the 222 nm emission spectrum. A deuterium lamp standard with a NIST-traceable spectral irradiance (Newport Model 63945, Irvine, CA) was used to radiometrically calibrate the UV spectrometer.

### Far-UVC dosimetry

Optical power measurements were performed using an 818-UV/DB low-power UV enhanced silicon photodetector with an 843-R optical power meter (Newport, Irvine, CA). Additional dosimetry to determine the uniformity of the UV exposure was performed using far-UVC sensitive film as described in our previous work^28,29^. This film has a high spatial resolution with the ability to resolve features to at least 25 μm, and exhibits a nearly ideal cosine response^30,31^. Measurements were taken between experiments therefore allowing placement of sensors inside the chamber.

A range of far-UVC exposures, from 3.6 μJ/cm^2^ up to 281.6 mJ/cm^2^, were used to define a response calibration curve. Films were scanned as 48 bit RGB TIFF images at 150 dpi using an Epson Perfection V700 Photo flatbed scanner (Epson, Japan) and analyzed with radiochromic film analysis software^32^ to calculate the total exposure based on measured changes in optical density.

Measurements using both a silicon detector and UV sensitive films were combined to compute the total dose received by a particle traversing the exposure window. The three vertically stacked lamps produced a nearly uniform dose distribution along the vertical axis thus every particle passing horizontally through the irradiation chamber received an identical dose. The lamp width (100 mm) was smaller than the width of the irradiation chamber window (260 mm) so the lamp power was higher near the center of the irradiation chamber window compared to the edge. The UV sensitive film indicated a power of approximately 120 μW/cm^2^ in the center third of the window and 70 μW/cm^2^ for the outer thirds. The silicon detector was used to quantify the reflectivity of the aluminum sheet at approximately 15% of the incident power. Combining this data allowed the calculation of the average total dose of 2.0 mJ/cm^2^ to a particle traversing the window in 20 seconds. Additionally, the silicon detector was used to confirm the attenuation of 222-nm light through a single sheet of plastic film was 65%. The addition of one or two sheets of plastic film between the lamps and the irradiation chamber window yielded average doses of 1.3 mJ/cm^2^ and 0.8 mJ/cm^2^, respectively.

### Benchtop aerosol irradiation chamber

A one-pass, dynamic aerosol / virus irradiation chamber was constructed in a similar configuration to that used by Ko et al.^33^, Lai et al.^34^, and McDevitt et al.^19,35^. A schematic overview of the system is shown in Fig. 3. Aerosolized viruses were generated by adding a virus solution into a high-output extended aerosol respiratory therapy (HEART) nebulizer (Westmed, Tucson, AZ) and operated using a dual-head pump (Thermo Fisher 420–2901–00FK, Waltham, MA) with an input flow rate of 11 L/min. The aerosolized virus flowed into the irradiation chamber where it was mixed with independently controlled inputs of humidified and dried air. Humidified air was produced by bubbling air through water, while dry air was provided by passing air through a desiccant air dryer (X06–02–00, Wilkerson Corp, Richland, MI). Adjusting the ratio of humid and dry air enabled control of the relative humidity (RH) within the irradiation chamber which, along with the nebulizer settings, determined the aerosol particle size distribution. An optimal RH value of 55% resulted in a distribution of aerosol particle sizes similar to the natural distribution from human coughing and breathing, which has been shown to be distributed around approximately 1 μm, with a significant tail of particles less than 1 μm^36–38^.

**Figure 3.**
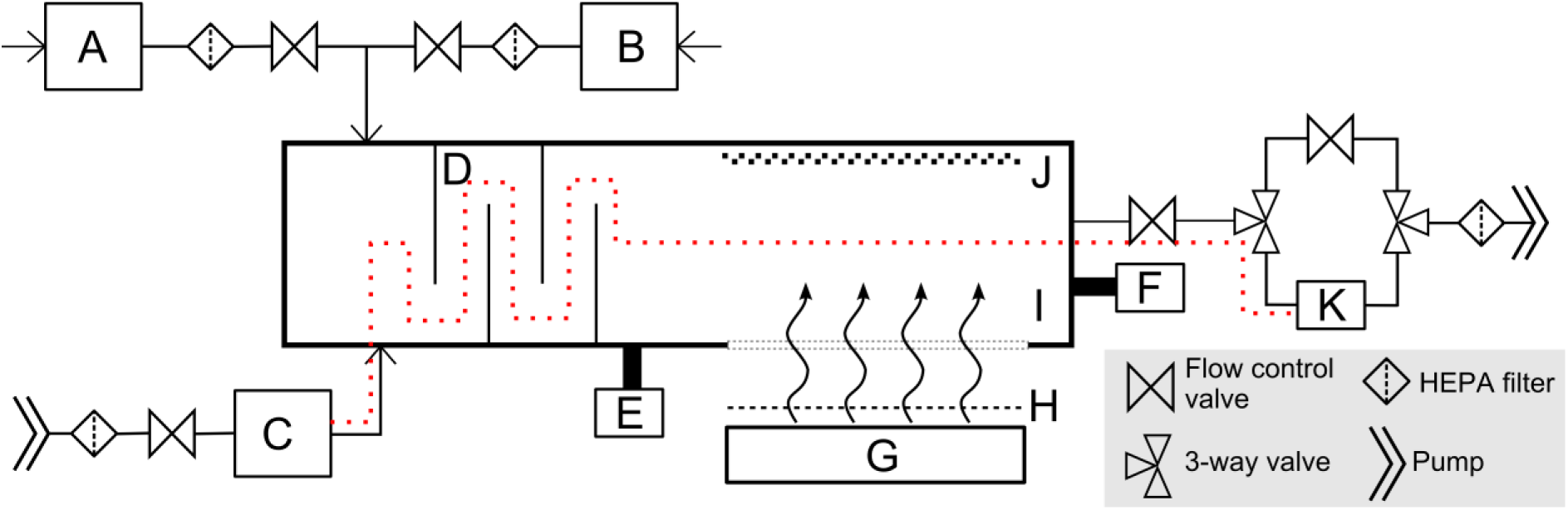
Schematic diagram of the custom UV irradiation chamber. The chamber is depicted in a top down view. Components of the setup include: water bubbler for humidified air input (A), a desiccator for dry air input (B), a nebulizer (C), baffles (D), an RH and temperature meter (E), a particle sizer (F), far-UVC lamps (G), band pass filters (H), a far-UVC transmitting plastic window (I), a reflective aluminum surface (J), and a BioSampler (K). Pumps are used to pressurize the nebulizer for aerosol generation and to control flow through the system. Flow control valves allow adjustments through the system. HEPA filters are included on all air inputs and outputs. A set of three way valves controls flow to or around the BioSampler. The vertically stacked lamps are directed at the window in the side of the chamber to expose the aerosols passing horizontally. The additional films to uniformly decrease the dose were placed between the filters and the window. The path of the aerosolized virus within the system during sampling is indicated with the red dotted line.

After combining the humidity control inputs with the aerosolized virus, input flow was directed through a series of baffles that promoted droplet drying and mixing to produce an even particle distribution ^34^. The RH and temperature inside the irradiation chamber were monitored using an Omega RH32 meter (Omega Engineering Inc., Stamford, CT) immediately following the baffles. A Hal Technologies HAL-HPC300 particle sizer (Fontana, CA) was adjoined to the irradiation chamber to allow for sampling of particle sizes throughout operation.

During UV exposure, the three 222-nm lamps with filters were stacked vertically and placed 11 cm from the irradiation chamber window. The lamps were directed at the 26 cm × 25.6 cm chamber window which was constructed of 254-μm thick UV transparent plastic film (Topas 8007X10, Topas Advanced Polymers, Florence, KY), and which had a transmission of ~65% at 222 nm. The wall of the irradiation chamber opposite the transparent window was constructed with polished aluminum in order to reflect a portion of the UVC light back through the exposure region, therefore increasing the overall exposure dose by having photons pass in both directions. The depth of the irradiation chamber between the window and the aluminum panel was 6.3 cm, creating a total exposure volume of 4.2 L.

Flow of the aerosols continues out of the irradiation chamber to a set of three way valves that could be configured to either pass through a bypass channel (used when no sampling was required), or a BioSampler (SKC Inc, Eighty Four, PA) used to collect the virus. The BioSampler uses sonic flow impingement upon a liquid surface to collect aerosols when operated at an air flow of 12.5 L/min. Finally, flow continued out of the system through a final HEPA filter and to a vacuum pump (WP6111560, EMD Millipore, Billerica, MA). The vacuum pump at the end of the system powered flow through the irradiation chamber. The flow rate through the system was governed by the BioSampler. Given the flow rate and the total exposure volume of the irradiation chamber, 4.2 L, a single aerosol droplet passed through the exposure volume in approximately 20 seconds.

The entire irradiation chamber was set up inside a certified class II type A2 biosafety cabinet (Labconco, Kansas City, MO). All air inputs and outputs were equipped with HEPA filters (GE Healthcare Bio-Sciences, Pittsburgh, PA) to prevent unwanted contamination from entering the chamber as well as to block any of the virus from releasing into the environment.

### Irradiation chamber performance

The custom irradiation chamber simulated the transmission of aerosolized viruses produced via human coughing and breathing. The chamber operated at a relative humidity of 55% which resulted in a particle size distribution of 87% between 0.3 μm and 0.5 μm, 11% between 0.5 μm and 0.7 μm, and 2% > 0.7 μm. Aerosolized viruses were efficiently transmitted through the system as evidenced from the control (zero exposure) showing clear virus integration (Fig. 1, top left).

### Experimental protocol

The virus solution in the nebulizer consisted of 1 ml of Dulbecco’s Modified Eagle’s Medium (DMEM, Life Technologies, Grand Island, NY) containing 10^8^ focus forming units per ml (FFU/ml) of influenza A virus [A/PR/8/34 (H1N1)], 20 ml of deionized water, and 0.05 ml of Hank’s Balanced Salt Solution with calcium and magnesium (HBSS^++^). The irradiation chamber was operated with aerosolized virus particles flowing through the chamber and the bypass channel for 15 minutes prior to sampling, in order to establish the desired RH value of ~55%. Sample collection initiated by changing air flow from the bypass channel to the BioSampler using the set of three way valves. The BioSampler was initially filled with 20 ml of HBSS^++^ to capture the aerosol. During each sampling time, which lasted for 30 minutes, the inside of the irradiation chamber was exposed to 222 nm far-UVC light through the UVC semi-transparent plastic window. Variation of the far-UVC dose delivered to aerosol particles was achieved by inserting additional UVC semi-transparent plastic films, identical to the material used as the chamber window, between the lamps and the chamber window. The extra plastic films uniformly reduced the power entering the chamber. The three test doses of 0.8, 1.3 and 2.0 mJ/cm^2^, were achieved by adding two, one, or no additional plastic films, respectively. Zero-dose control studies were conducted with the excimer lamps turned off. Experiments at each dose were repeated in triplicate. After the sampling period was completed the solution from the BioSampler was used for the virus infectivity assay.

### Virus infectivity assay

We measured viral infectivity with a focus forming assay that employs standard fluorescent immunostaining techniques to detect infected host cells and infectious virus particles^39^. Briefly, after running through the irradiation chamber for 30 minutes, 0.5 ml of virus suspension collected from the BioSampler was overlaid on a monolayer of Madin-Darby Canine Kidney (MDCK) epithelial cells routinely grown in DMEM supplemented with 10% Fetal Bovine Serum (FBS), 2 mM L-alanyl-L-glutamine, 100 U/ml penicillin and 100 μg/ml streptomycin (Sigma-Aldrich Corp. St. Louis, MO, USA). Cells were incubated with the virus for 45 minutes, washed three times with HBSS^++^ and incubated overnight in DMEM. Infected cells were then fixed in 100% ice cold methanol at 4°C for 5 minutes and labeled with influenza A virus nucleoprotein antibody [C43] (Abcam ab128193, Cambridge, MA) 1:200 in HBSS^++^ containing 1% bovine serum albumin (BSA; Sigma-Aldrich Corp. St. Louis, MO, USA) at room temperature for 30 minutes with gentle shaking. Cells were washed three times in HBSS^++^ and labeled with goat anti-mouse Alexa Fluor-488 (Life Technologies, Grand Island, NY) 1:800 in HBSS^++^ containing 1% BSA at room temperature for 30 minutes with gentle shaking. Following three washes in HBSS^++^, the cells were stained with Vectashield containing DAPI (4’,6-diamidino-2-phenylindole) (Victor Laboratories, Burlingame, CA) and observed with the 10x and 40x objectives of an Olympus IX70 fluorescent microscope equipped with a Photometrics PVCAM high-resolution, high-efficiency digital camera. For each sample, at least three fields of view of merged DAPI and Alexa-488 images were acquired. Image-Pro Plus 6.0 software (Media Cybernetics, Bethesda, MD) was used to analyze the 10x images to measure the FFUuv as the ratio of cells infected with the virus divided by the total number of cells.

### Data analysis

The surviving fraction (*S*) of the virus was calculated by dividing the fraction of cells that yielded positive virus growth at each UV dose (FFU_uv_) by the fraction at zero dose (FFU_controls_): *S*=FFU_uv_/FFU_controls_. Survival values were calculated for each repeat experiment and natural log (ln) transformed to bring the error distribution closer to normal^40^. Linear regression was performed using these normalized ln[*S*] values as the dependent variable and UV dose (D, mJ/cm^2^) as the independent variable. Using this approach, the virus survival (*S*) was fitted to first-order kinetics according to the equation^7^:

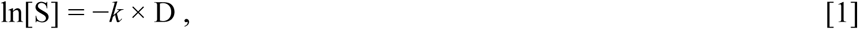

where *k* is the UV inactivation rate constant or susceptibility factor (cm^2^/mJ). The regression was performed with the intercept term set to zero, which represents the definition of 100% relative survival at zero UV dose. Bootstrap 95% confidence intervals for the parameter *k* were calculated using R 3.2.3 software^41^. The virus inactivation cross section, D_95_, which is the UV dose that inactivates 95% of the exposed virus, was calculated as D_95_ = − ln[1 − 0.95]/*k*.

## Acknowledgements

This work was supported by the Shostack Foundation and also NIH grant 1R41AI125006–01. We thank Dr. Rea Dabelic from the Department of Environmental Health Sciences, Mailman School of Public Health at Columbia University for her expertise and training with viral cell culture.

## Author Contributions

D.W., M.B., and V.G. designed and performed experiments, analyzed the data, and wrote the manuscript; I.S. analyzed the data; C.C. and A.W.B. designed the irradiation chamber; G.W.J. constructed the irradiation chamber; D.J.B and G.R-P. supervised, contributed conceptual advice, and wrote the manuscript. All authors discussed the results and commented on the manuscript.

